# Collisional cross-section prediction for multiconformational peptide ions with IM2Deep

**DOI:** 10.1101/2025.02.18.638865

**Authors:** Robbe Devreese, Alireza Nameni, Arthur Declercq, Emmy Terryn, Ralf Gabriels, Francis Impens, Kris Gevaert, Lennart Martens, Robbin Bouwmeester

**Affiliations:** VIB-UGent Center for Medical Biotechnology, VIB, Ghent, Belgium; Department of Biomolecular Medicine, Ghent University, Ghent, Belgium; BioOrganic Mass Spectrometry Laboratory (LSMBO), IPHC UMR 7178, University of Strasbourg, CNRS, ProFI FR2048, Strasbourg, France

## Abstract

Peptide collisional cross-section (CCS) prediction is complicated by the tendency of peptide ions to exhibit multiple conformations in the gas phase. This adds further complexity to downstream analysis of proteomics data, for example for identification or quantification through feature finding. Here, we present an improved version of IM2Deep that is trained on a carefully curated dataset to predict CCS values of multiconformational peptides. The training data is derived from a large and comprehensive set of publicly available datasets. This comprehensive training dataset together with a tailored architecture allows for the accurate CCS prediction of multiple peptide conformational states. Furthermore, the enhanced IM2Deep model also retains high precision for peptides with a single observed conformation. IM2Deep is publicly available under a permissive open source license at https://github.com/compomics/IM2Deep.

## Introduction

In recent years, technical advances in LC-IM-MS/MS technology have strongly enhanced identification capabilities in complex proteomics workflows (1–3). Traditionally, LC-MS/MS systems solely depended on liquid chromatography and mass analyzers for peptide separation and selection respectively, before fragmentation. In contrast, MS instruments incorporating ion mobility, like the timsTOF series, allow for gas-phase ion separation with ion mobility spectrometry (IMS) and use parallel accumulation–serial fragmentation (PASEF) technology to increase sensitivity and acquisition speed (4). PASEF allows precursor ions to accumulate in the trapped ion mobility spectrometry (TIMS) tunnel before being sequentially released, a feature particularly advantageous for increasing the overall sensitivity for low-abundant ions and lowering spectrum complexity.

The inversed reduced ion mobility (1/K0) and the mass-to-charge ratio of a peptide, measured by IMS and MS respectively, can be used to calculate the collisional cross section (CCS) of said peptide ion using the Mason-Schamp equation (5). CCS represents the rotationally averaged effective area with which an ion collides with a neutral gas. It is closely tied to the ion’s chemical structure and three-dimensional conformation, making it an important distinguishing characteristic of an ion in the gas phase. This characteristic is useful for improved identification confidence upon comparing predicted and observed CCS values (6,7). Furthermore, a predicted CCS value can also be used for more accurate quantification through feature finding (8) and to prioritize acquisition time (9). The potential benefit from predicted CCS values led to the development of various machine learning models based on amino acid- or atom-level features and physicochemical properties (6,7,10). However, current machine learning-based CCS prediction models overlook the possibility of peptide ions adopting multiple conformations in the gas phase (Figure 1), as these are often trained on the most abundant conformer within a dataset. This approach can result in the exclusion of valuable data and biases the model towards the most prevalent conformers in each experiment, inherently assuming that the dominant conformer in one experiment will consistently dominate in others, which is not a foregone conclusion and might result in inaccurate prediction.

**Figure 1:**
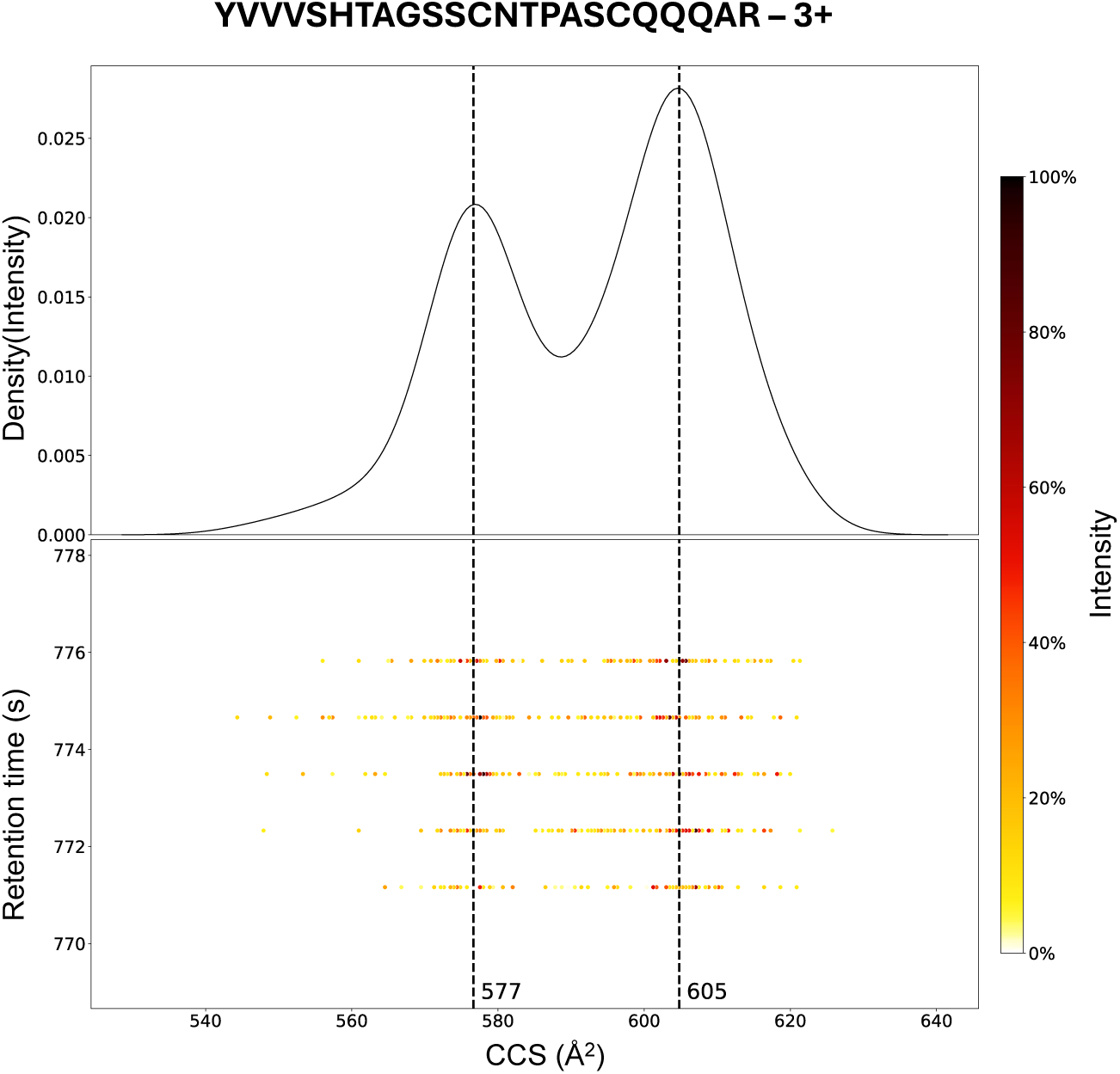
Ion mobility distribution of a representative peptide precursor ion (UniProt ID of inferred protein: O75594, data from PRIDE accession PXD046507). The upper panel shows the ion mobility distribution with multiple distinct peaks, indicating the presence of multiple conformations. The lower panel displays the ion intensity plotted against retention time (s) and CCS (Å²). MaxQuant reported two CCS features for this precursor, one at 577 Å² and one at 605 Å².

Previous studies have identified several factors contributing to the multiconformationality of precursor ions in LC-IM-MS/MS experiments. The emergence and distribution of these conformers can depend on the composition of the solvent from which the peptide is electrosprayed (11), suggesting that different conformational states in solution can produce distinct gas-phase conformers upon dehydration, that are distinguishable by IMS-MS. Additionally, peptide ions may adopt varying conformations depending on the activation voltage applied (12), and molecular modeling indicates that *cis-trans* isomerization of proline residues may also play a role in influencing conformation (13). Furthermore, charge localization within the peptide when there is no preferential charge site can also result in multiple gas phase conformations (14).

The rising popularity of ion mobility-assisted LC-MS/MS experiments, coupled with an increasing commitment to open science within the proteomics community, has led to a significant expansion in publicly available TIMS-acquired proteomics datasets. These datasets are critical for advancing data-driven proteomics software. Deep learning models, which rely on large volumes of data for accurate predictions, particularly benefit from such datasets. By pooling multiple datasets, more comprehensive training sets can be created, especially for less common phenomena like multiconformational peptide ion detection. While a single LC-IM-MS/MS dataset might not yield enough training data for accurate multiconformer CCS predictions, aggregating datasets is expected to enhance both the quantity and reliability of training samples.

We have recently introduced IM2Deep, a deep learning-based peptide CCS predictor that builds on the principles of DeepLC, a retention time predictor (7,15). IM2Deep encodes peptidoforms at the atomic composition level, enabling precise CCS predictions for peptides with modifications not encountered during training. In this study, we present an improved version of IM2Deep designed to predict CCS values for peptide ions exhibiting multiple conformations in the gas phase. This was achieved with a modified architecture of IM2Deep to support multi-output predictions together with transfer learning to tailor the model for a much smaller dataset of multiconformational peptide ions. These changes to the training data and architecture enable IM2Deep to accurately predict CCS values for multiconformational peptides.

## Methods

### Datasets

We searched the PRIDE public proteomics data repository for datasets generated using timsTOF Pro and timsTOF Pro 2 instruments, focusing on those with readily available identification output, including CCS annotations. In total, we collected MaxQuant (16) evidence output from 30 different PRIDE projects, totaling 1,248 LC-IM-MS/MS runs for further processing (Supplementary Table 1).

### Creation of a multiconformer peptide dataset

This section details the steps taken to aggregate the separate experimental datasets into a unified dataset of multiconformational peptides (Supplementary Figure S1). First of all, only peptide ions identified by separate MS1 clusters (features) were kept for further processing. The initial step involved aligning CCS values across all LC-IM-MS/MS runs in each experiment. This was done by calculating a charge-specific linear offset (y = x + b) between overlapping peptide-charge state pairs, similar to the method described by Meier *et al*. (10). For each peptide-charge state pair, the CCS of the identification with the highest intensity was used as the alignment reference, but CCS values of the less abundant identifications were also aligned and retained. Beginning with the run containing the highest number of identifications, all other runs were sequentially aligned and added to the dataset in descending order of identifications. To ensure good alignment, runs with less than 100 overlapping peptide-charge state pairs with the aligned dataset (n = 284) were ignored.

Following CCS alignment, conformers were identified within each LC-IM-MS/MS run. Identical precursor identifications exhibiting a larger than 2% difference in CCS within the same run were considered distinct conformational states, as suggested by previous research (10).

To increase the confidence in accurately identifying conformers, we filtered the dataset by checking for recurring conformers across multiple separate runs. Specifically, we compared the CCS values reported for each precursor in each run. Only multiconformational precursors with multiple CCS values that could be matched to corresponding values in other runs, within a 2% tolerance, were retained. The mean CCS value for each conformer was retained in the dataset.

### Creation of a uniconformer peptide dataset

Precursors that did not meet the above criteria were classified as uniconformational and placed in a separate dataset. This dataset underwent additional filtering to confirm that multiconformational precursors, where distinct conformers appeared only between runs (and not within the same run), were excluded. Thus, only precursors with CCS values showing no deviation greater than 2% between aligned runs were included in the uniconformer peptide dataset.

### IM2Deep architecture for multiconformer CCS prediction

The architecture of IM2Deep as described in Declercq & Devreese *et al*. (7) is nearly identical as the one of DeepLC (15). Briefly, IM2Deep employs a convolutional architecture with four distinct pathways through which each encoded peptide is processed. Three of these pathways employ convolutional and maximum pooling layers to capture local structures, handling atomic composition of amino acids, diamino acids, and one-hot encoding for unmodified amino acids. A fourth pathway processes global features using densely connected layers. The outputs from all paths are combined, flattened, and passed through six dense layers in the final combined path, culminating in a single output node that predicts CCS.

To enable multiconformer CCS prediction, this architecture was adapted to support multi-output prediction. Instead of a single output node, IM2Deep now features a branched output with two final dense layers, each ending in a single output node, producing a distinct CCS prediction. We trained the multi-output IM2Deep models both with and without a pretrained single-output model as foundation. The single-output model had been trained on the dataset used in Declercq & Devreese *et al*. (7). Pretrained model weights were loaded into the shared layers of the multi-output model prior to training, with all parameters remaining fully trainable.

The multi-output IM2Deep models were trained and evaluated on the multiconformer dataset, which was randomly split into 81% training, 9% validation, and 10% testing sets. Training continued for a maximum of 500 epochs, with early stopping based on validation set performance to prevent overfitting.

To ensure distinct predictions for each target, we implemented a custom loss function. Simple loss functions, such as the sum of mean absolute errors (MAEs) between each prediction-target pair, can lead to cases where both predictions converge to the same target. Our custom loss function was designed to address this issue by forcing the model to produce two distinct predictions corresponding to two separate targets. Firstly, both targets and model outputs are ordered by size to impose a consistent prediction-target pairing.

The mean absolute error is calculated between each ordered prediction and its corresponding ordered target:

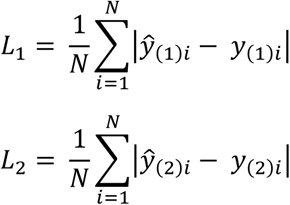

where N is batch size and

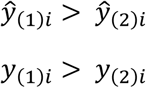

Additionally, the difference between the targets and the predictions is computed:

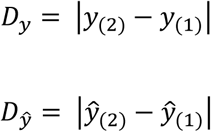

The mean absolute error of these differences is then calculated:

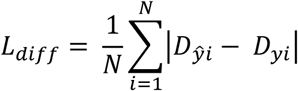

The total loss is then calculated as the sum of the two target-prediction MAEs and the MAE of the differences, multiplied by a weight to emphasize the importance of the difference term:

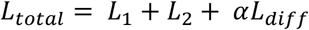

This custom loss function ensures that each prediction corresponds closely to a distinct target. For the final models, this difference weight was set to five, as determined by achieving the highest accuracy on the validation set in a tuning experiment where the weight was set from zero up to ten (Supplementary Figure S2).

### Visualization of ion mobilograms

To illustrate the advantages of the multiconformational IM2Deep model, ion mobilograms were generated to visualize ion mobility distributions for identified precursors from an independent public LC-IM-MS/MS dataset (PXD046507) (17). The raw timsTOF measurements were accessed from the raw .d folder using the Alphatims (version 1.0.8) Python package (18). Precursors identified by MaxQuant were matched to the raw ion mobility measurements based on their retention time and *m/z* values, with a retention time tolerance of 4 s and a *m/z* precursor tolerance of 10 ppm. Ion mobility measurements were converted to CCS using the Mason-Schamp equation (5).

## Results

### Description of the (multi)conformer dataset

In this study, we compiled identification data from 1,248 LC-IM-MS/MS runs across 30 PRIDE projects to determine peptide conformers. To enhance the reliability of our dataset, we excluded conformers not identified in multiple runs (Supplementary Figure S3).

After filtering the dataset, the final multiconformer dataset included 29,691 unique peptidoform-charge state pairs. This is about 4.1% the size of the uniconformer dataset (n = 728,249), which is in line with previously published estimates (6,10). The number of observed conformational states for a single precursor in the multiconformer dataset reached as high as five (n = 5, Supplementary Figure 4).

We identified differences in the physicochemical properties of uni- and multiconformational subsets of precursor ions (Figure 2). Notably, multiconformational precursor ions tended to be longer (Figure 2A) and carried higher charges (Figure 2B). These observations support findings by Meier *et al*., who reported that prediction models trained to estimate only one CCS value show a bias, particularly affecting the accuracy for longer, 3+ and 4+ charged peptides (10).

**Figure 2.**
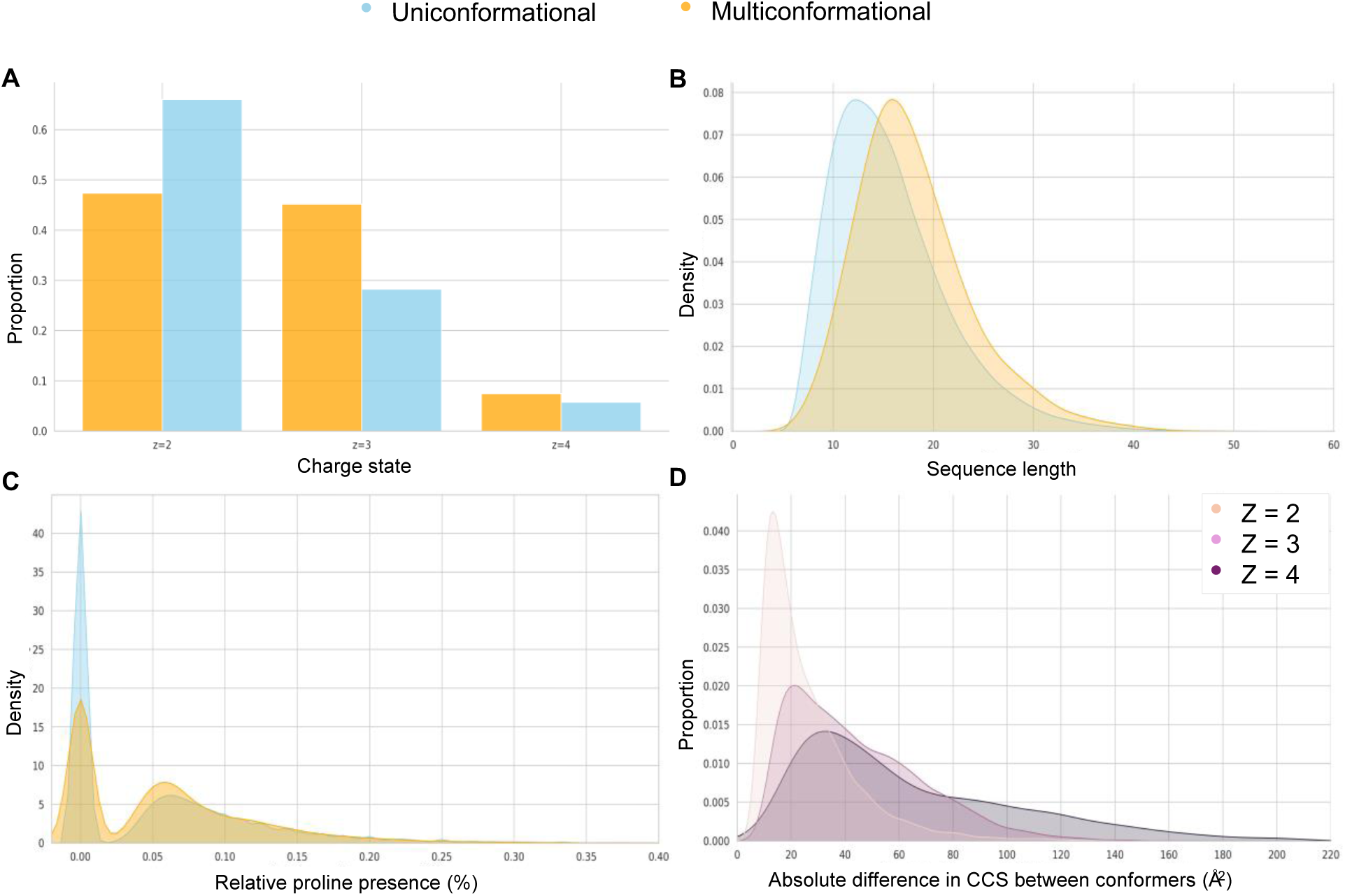
Physicochemical properties and CCS differences between uniconformational (blue) and multiconformational (orange) peptides. A) Distribution of charge states. B) Sequence length distribution. C) Relative frequency of proline residues. D) Distribution of absolute differences in CCS between conformers for peptides with different charge states.

Given the hypothesis that *cis*-*trans* isomerization of proline residues could contribute to multiconformationality, we examined the presence of proline within the two subsets. Indeed, we identified an enrichment in proline presence within the multiconformational peptides (61.9%), compared to peptides not exhibiting multiple conformations (52.4%). Additionally, the relative frequency of proline residues-defined as the number of prolines divided by the length of the sequence-is also higher in the multiconformer subset (Figure 2C). Furthermore, peptides with higher charge states exhibited greater differences in CCS between conformers (Figure 2D), with a broader distribution of CCS differences observed among highly charged peptide ions. Small amino acids, such as alanine and valine also occur more often in the multiconformational peptides, while longer amino acids such as glutamic acid, lysine and arginine had a lower relative presence (Supplementary Figure S5).

### Multi-output IM2Deep for two conformer CCS prediction

Using the multiconformer dataset, filtered for precursors exhibiting two conformations (n = 26,970), we trained a new version of the IM2Deep model specifically adapted for multi-output prediction, targeting two CCS values corresponding to each conformer. We employed two training strategies: one involved training a model from scratch, while the other utilized fine-tuning. In the fine-tuning approach, weights from a pre-trained single-output IM2Deep model were loaded into the shared layers of the multi-output model before training started. This method leverages the features learned from the pre-trained model to enhance prediction accuracy, a strategy that has previously demonstrated success in challenging peptide property predictions (19,20).

Learning curves on the validation set reveal that the fine-tuned model outperforms the newly trained model (Figure 3). The test set results further corroborate these findings (Figure 4), with the fine-tuned model showing superior prediction accuracy compared to the model trained with random initialization. Importantly, the median relative error for both CCS predictions in the fine-tuned model remains well below the 2% threshold used to define multiconformationality. This suggests that the model can effectively distinguish between different conformers and accurately predicts their corresponding CCS values with high precision. While both training and testing datasets contained predominantly tryptic peptides, the model’s performance on non-tryptic peptides is comparable (Supplementary Figure S6).

**Figure 3:**
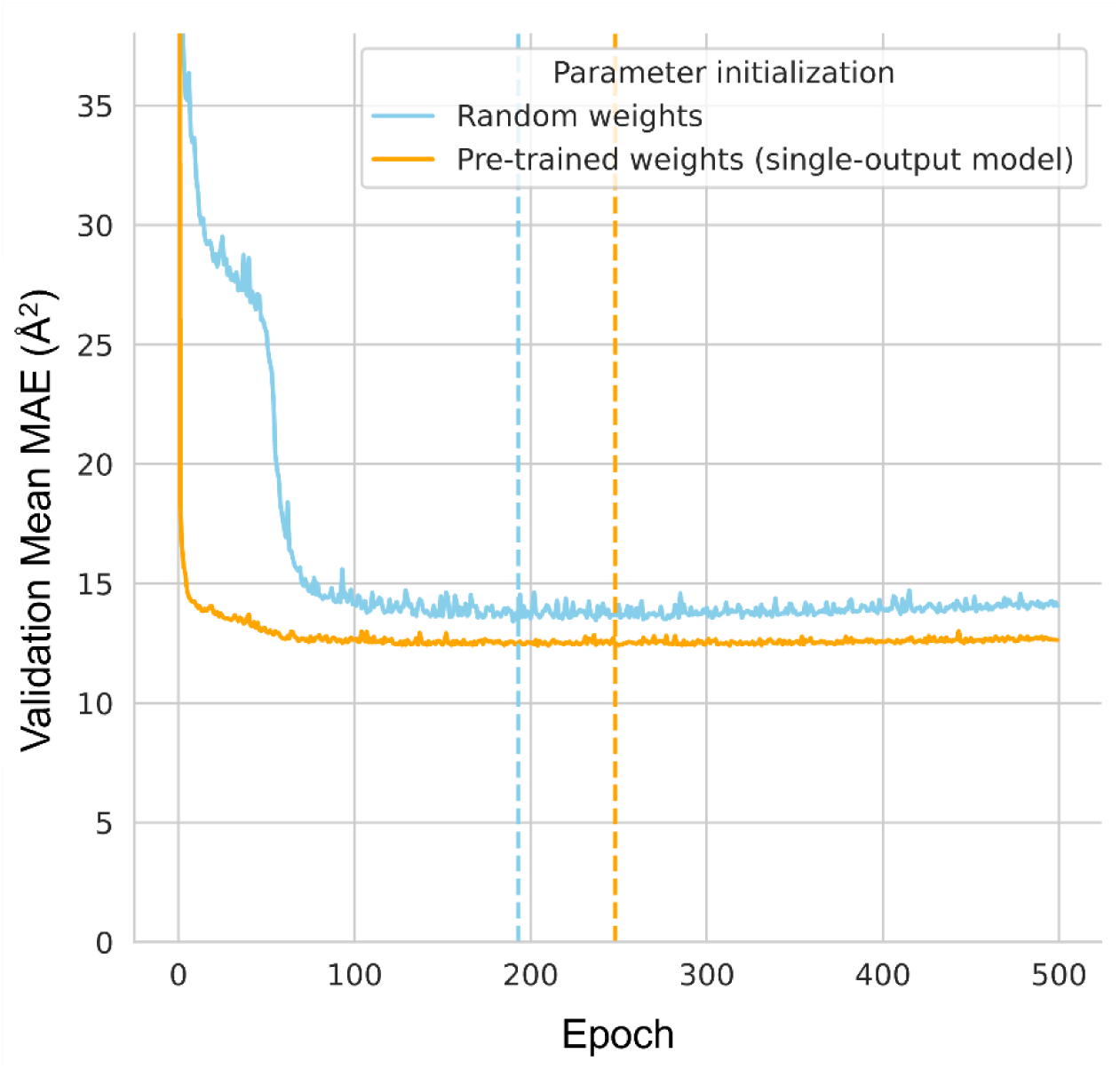
Performance curves over 500 training epochs for the multi-output IM2Deep model, comparing performance with different parameter initialization strategies. The model initialized with pre-trained weights from a single-output IM2Deep model (orange) shows consistently lower validation MAE compared to the model initialized with random weights (blue). Dotted lines represent the points at which the highest validation accuracy is achieved, i.e. where early stopping occurs.

**Figure 4:**
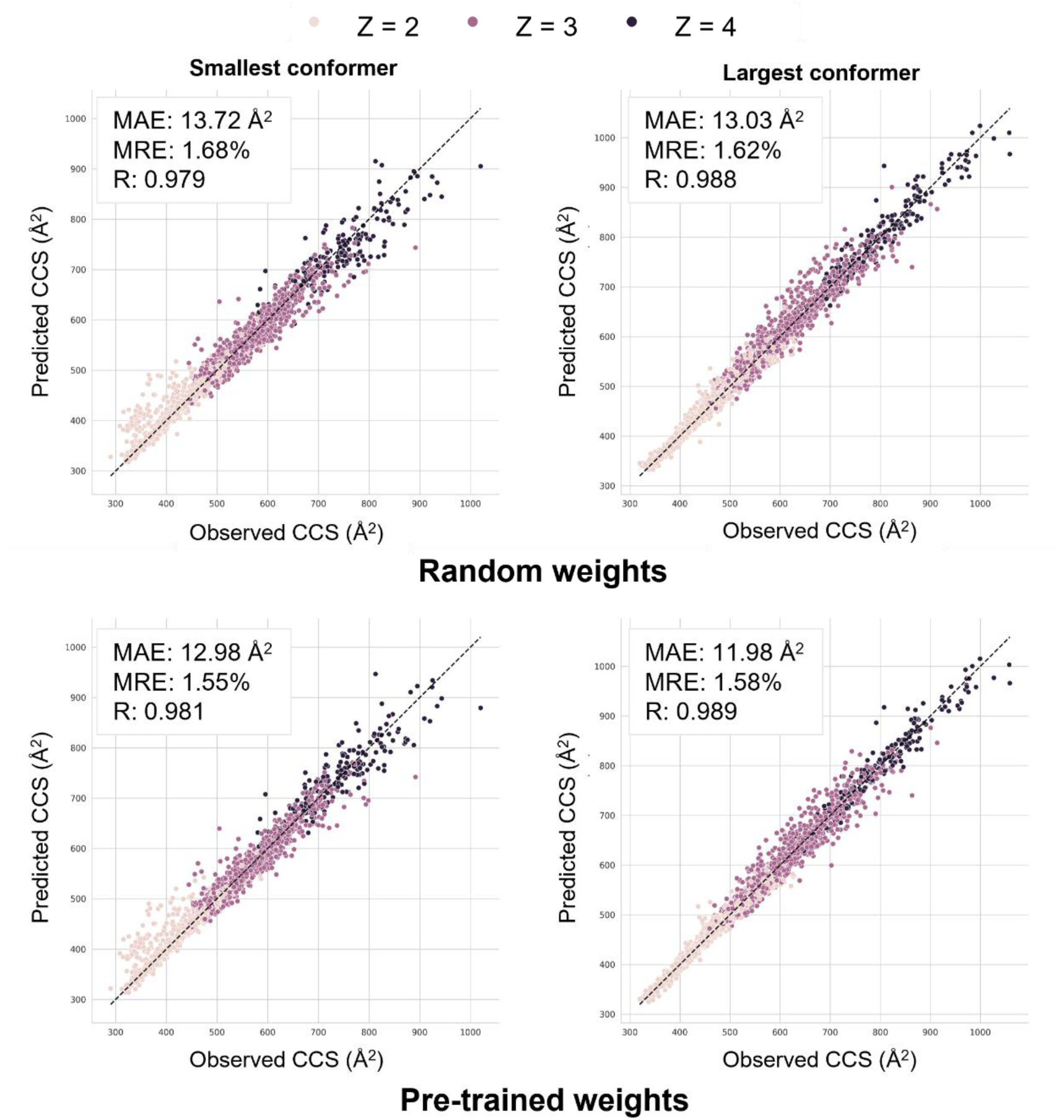
Scatter plots comparing predicted versus observed CCS values for the multi-output IM2Deep model trained with different parameter initialization strategies. The top row shows the performance of the model trained with random weight initialization, while the bottom row displays the performance of the model fine-tuned with pre-trained weights from a single-output IM2Deep model. Each point represents a peptide ion, with color indicating its charge state. The diagonal red line represents the ideal scenario where predicted CCS values match observed values. The left plots correspond to the first, smaller CCS prediction, and the right plots to the second, larger CCS prediction of each multiconformational precursor.

### Evaluating model robustness and comparison with baseline models

To ensure that an additional second prediction does not artificially improve its performance, a comparison is made with two baseline models: (1) a baseline model trained on CCS values from peptides exhibiting only one conformation (the uniconformer dataset, see methods); (2) a baseline model trained on the CCS values of one randomly selected conformation from each peptide in our multiconformational training set. Despite being trained on only one target CCS value per peptide, the architecture of these baseline models have two output nodes and thus allow for two predictions. These baseline models serve as benchmarks to evaluate the benefits of training a model on multiple CCS values for multiconformational predictions. The loss function used to train the baseline models is calculated as the sum of the mean absolute errors between each prediction and the target CCS.

As illustrated in Figure 5, the transfer-learned model strongly outperforms both baseline models in predicting multiple CCS values, with marked improvements in both MAE and Pearson correlation coefficients. These results underscore the model’s ability to effectively capture the multiconformational nature of these peptides, owing to its training on multiple conformations, and not just because of the capacity of the model to make two predictions per precursor.

**Figure 5:**
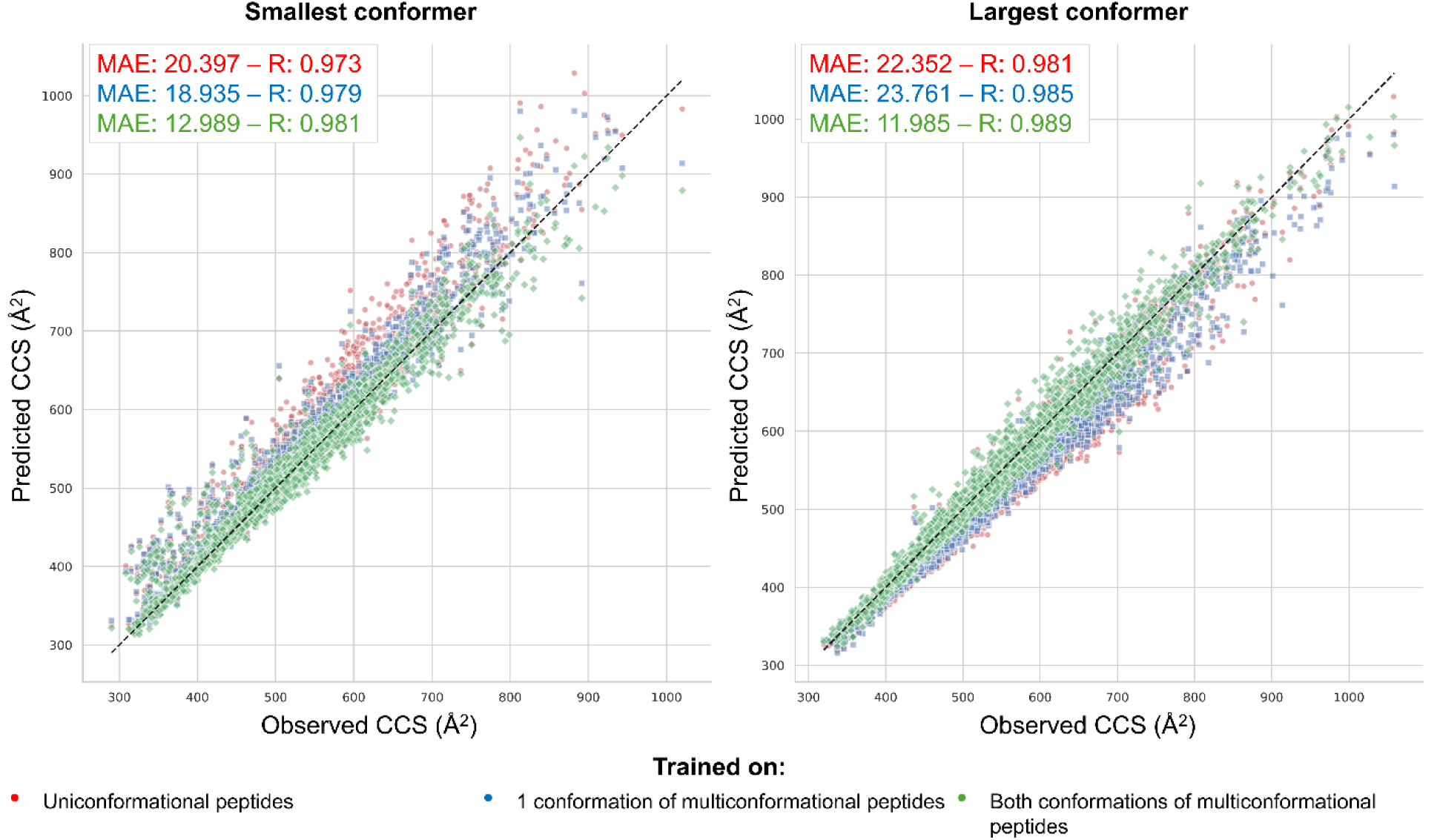
Scatter plots comparing predicted versus observed CCS values for the multi-output IM2Deep model and two baseline models, each trained on different datasets. The left plot shows the first CCS prediction, and the right plot shows the second CCS prediction for each peptide. Red dots represent predictions from the baseline model trained on peptides with only one conformation. Blue dots represent predictions from the baseline model trained on one randomly selected conformation from each multiconformational peptide. Green dots represent predictions from the transfer-learned multiconformational model trained on both conformations of multiconformational peptides. Each dot corresponds to a peptide ion, with the dashed line representing the ideal scenario where predicted CCS values match observed values.

An ideal multiconformational model should also accurately predict CCS values for peptides where only one conformation is observed. To assess this, we compared the accuracy of the closest prediction made by the transfer-learned multiconformational model to that of a model trained exclusively on uniconformational peptides, on a test set of uniconformational peptides. Although the multiconformational model has the advantage of making two predictions and selecting the best one, we controlled for this potential bias by allowing the baseline model trained on uniconformational peptides to also make two predictions, again selecting the best one.

The results, shown in Figure 6, indicate that the multiconformational model, despite being trained on peptides with multiple conformations, maintains or even improves its accuracy in predicting CCS values for peptides with only one observed conformation, particularly in terms of MAE and Pearson correlation. This improvement likely stems from the model’s ability to capture important structural information that is not fully represented in models trained on a single CCS value.

**Figure 6:**
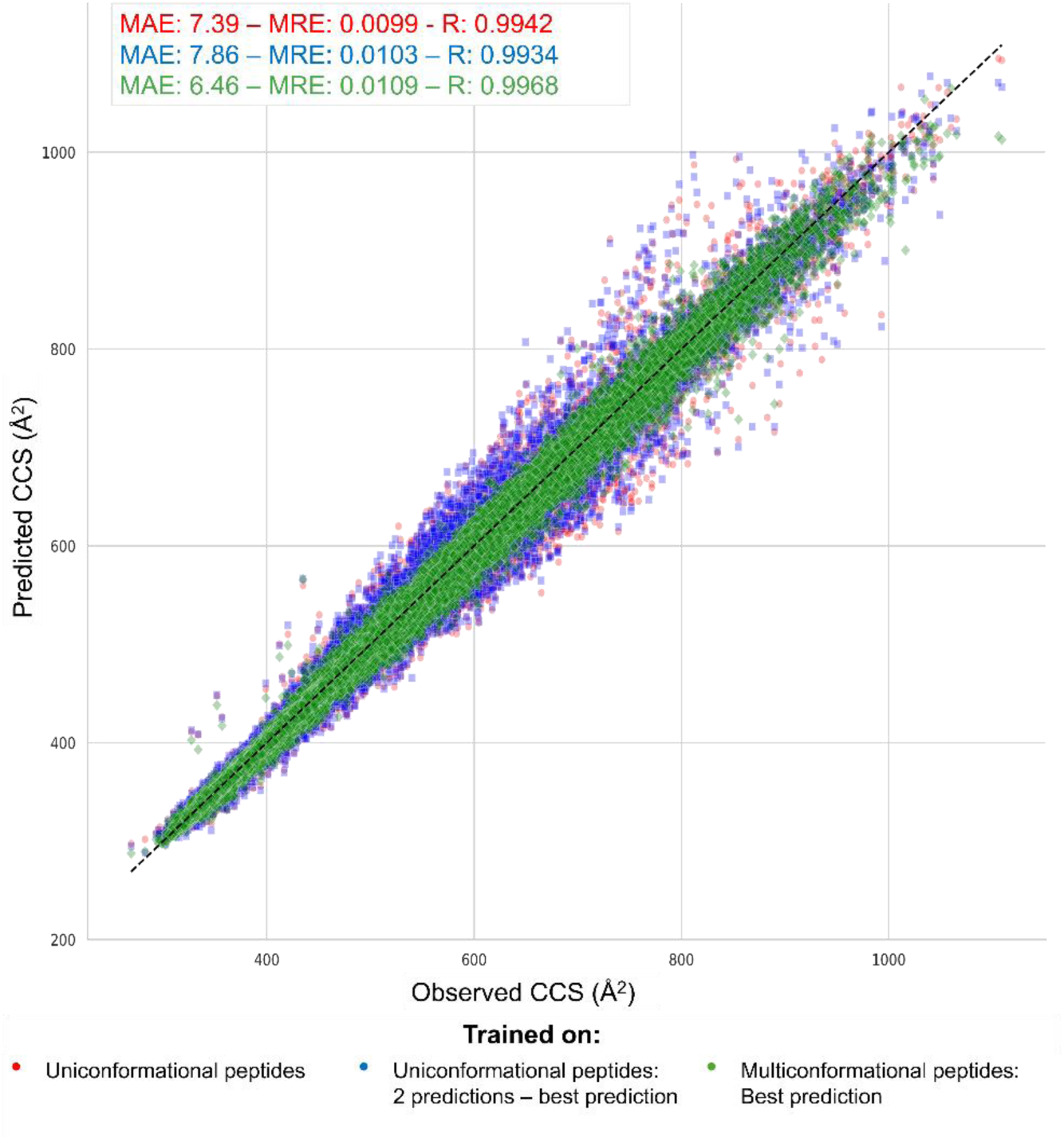
Scatter plot comparing predicted versus observed CCS values for uniconformational peptides using the different models.

### Examples demonstrating the advantages of the multiconformer model on an independent dataset

To showcase the capabilities of the multi-output IM2Deep model, we predicted two CCS values for all identified precursors from a run in a different public LC-IM-MS/MS experiment (PXD046507) and visualized these predictions on mobilograms derived from raw timsTOF data. This section highlights specific scenarios where the new multiconformational model offers added value over the original IM2Deep model, which was limited to predicting a single CCS value.

In Figure 7A, a precursor ion displays two distinct peaks in its ion mobility distribution, indicating the presence of two conformations. The original IM2Deep model successfully predicts the CCS for one of these conformers (red), while MaxQuant reports the CCS of the other, less intense conformer (black). In PSM rescoring, where the observed CCS values are compared to predicted ones, this discrepancy could lead to the penalization and potential exclusion of the PSM, even though IM2Deep made an accurate prediction. In contrast, the multiconformational model accurately predicts the CCS values for both conformations, eliminating this issue when the closest prediction to the observed CCS is used for rescoring.

**Figure 7:**
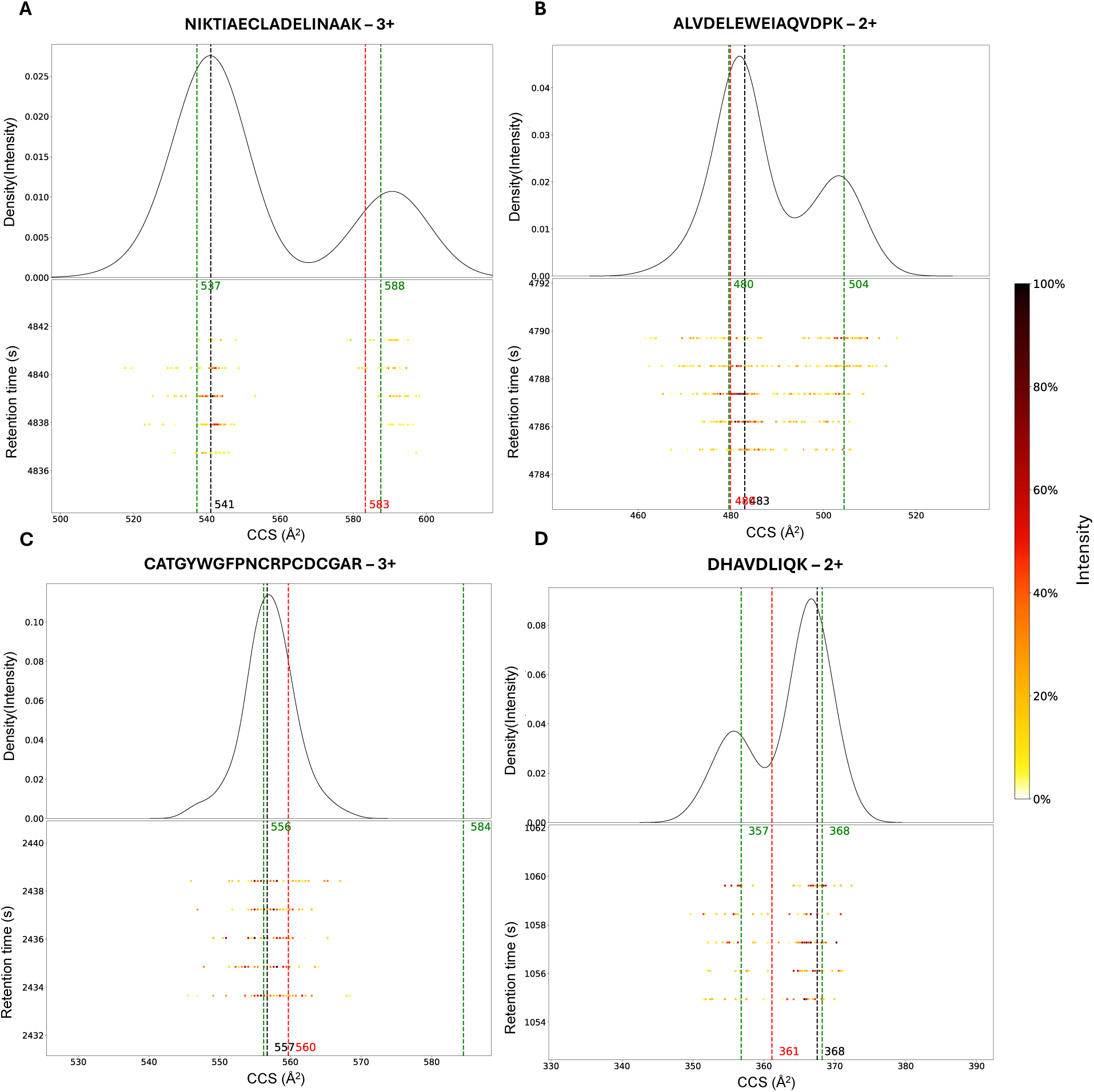
Mobilograms displaying the ion mobility distributions of selected precursor ions, highlighting the predictive capabilities of the multi-output IM2Deep model. CCS value reported by MaxQuant indicated by black dashed lines. Classical IM2Deep prediction (single-output) indicated by red dashed line. CCS predictions made by multiconformational model indicated by green dashed lines. A) Precursor ion (UniProt ID of inferred protein: P46782) showing two distinct peaks in the ion mobility distribution, where the original model predicts the CCS of the non-reported conformer. B) Precursor ion (UniProt ID of inferred protein: Q9Y2B0) showing two distinct conformations, where the multiconformational model accurately predicts CCS of a conformer not identified by MaxQuant. C) Precursor ion (UniProt ID of inferred protein: O15230) showing a single dominant conformation, accurately predicted by the multiconformational model. D) Precursor ion (UniProt IDs of inferred proteins: P0C0L4 and P0C0L5) where classical IM2Deep model prediction appears to be the average of two conformations. Observed and predicted CCS values are rounded to the nearest integer for simplicity.

An interesting advantage of the new multiconformational model is its ability to accurately predict CCS values for conformations not identified by the search engine. For example, as shown in Figure 7B, while MaxQuant reported the CCS value of the most intense conformer, the multiconformational model accurately predicted the CCS value of a less intense, unreported conformer. This capability offers deeper insights into the structural dynamics and conformational diversity of peptide ions in the gas phase.

In the majority of cases, peptide ions exhibit a single conformation (Figure 7C). As previously demonstrated, the multiconformational model still makes accurate predictions for these peptide ions. When the best prediction is selected, the second prediction might be redundant, but could also represent an accurate prediction of an unobserved conformer.

As discussed by others, a maximum likelihood estimation approach to CCS prediction typically converges to the mean CCS value of all conformers when no prior conformer filtering is performed, and a mean value for each peptide ion is retained in the final training set (6). Previous approaches have tried to avoid this by retaining only the most intense conformer in datasets to prevent erroneous convergence to the mean of all conformers. However, when datasets are combined, and CCS values are averaged over multiple datasets, this assumes that the most intense conformer of each peptide is consistent across all experiments. This assumption may not always hold. Indeed, there are instances where the original IM2Deep model, which relies on this assumption, predicts a CCS value close to the mean of two conformers (Figure 7D). In contrast, our new approach assumes this consistency only for calculating linear offsets to align CCS values between datasets, where the impact of uncommon multiconformational peptides should be minor. For training, however, no hierarchical order is imposed on conformers; instead, they are matched across datasets based on their CCS values, with an average value retained for each conformer. This approach allows for more accurate and flexible predictions, accounting for the possibility of multiple conformations not necessarily consistent between experiments.

It is important to note that none of the precursor ions used as examples in this analysis were included in the training data for the multiconformational model.

## Discussion

In this study, we developed a new multi-output IM2Deep model designed to predict CCS values for peptides that exhibit multiple conformations. Using a multiconformer dataset, generated by combining LC-IM-MS/MS data from various experiments and selecting peptides with multiple observed conformations, we trained the model to predict two distinct CCS values for each peptide. The results demonstrate that the multiconformational model accurately predicts CCS values for multiple conformers, while retaining its ability to make accurate predictions for peptides exhibiting only one conformation. This marks an important step forward in the field of CCS prediction.

In this study, we limited the training of the multi-output IM2Deep model to two CCS values due to the size limitations of our training set. While this approach provides important improvements, it is evident that predicting CCS values for higher-order conformations would further enhance performance for peptides showing more than two conformations (Figure 8B). Our current model architecture is flexible, requiring only slight adjustments to the architecture and training procedure to accommodate these higher-order predictions.

**Figure 8:**
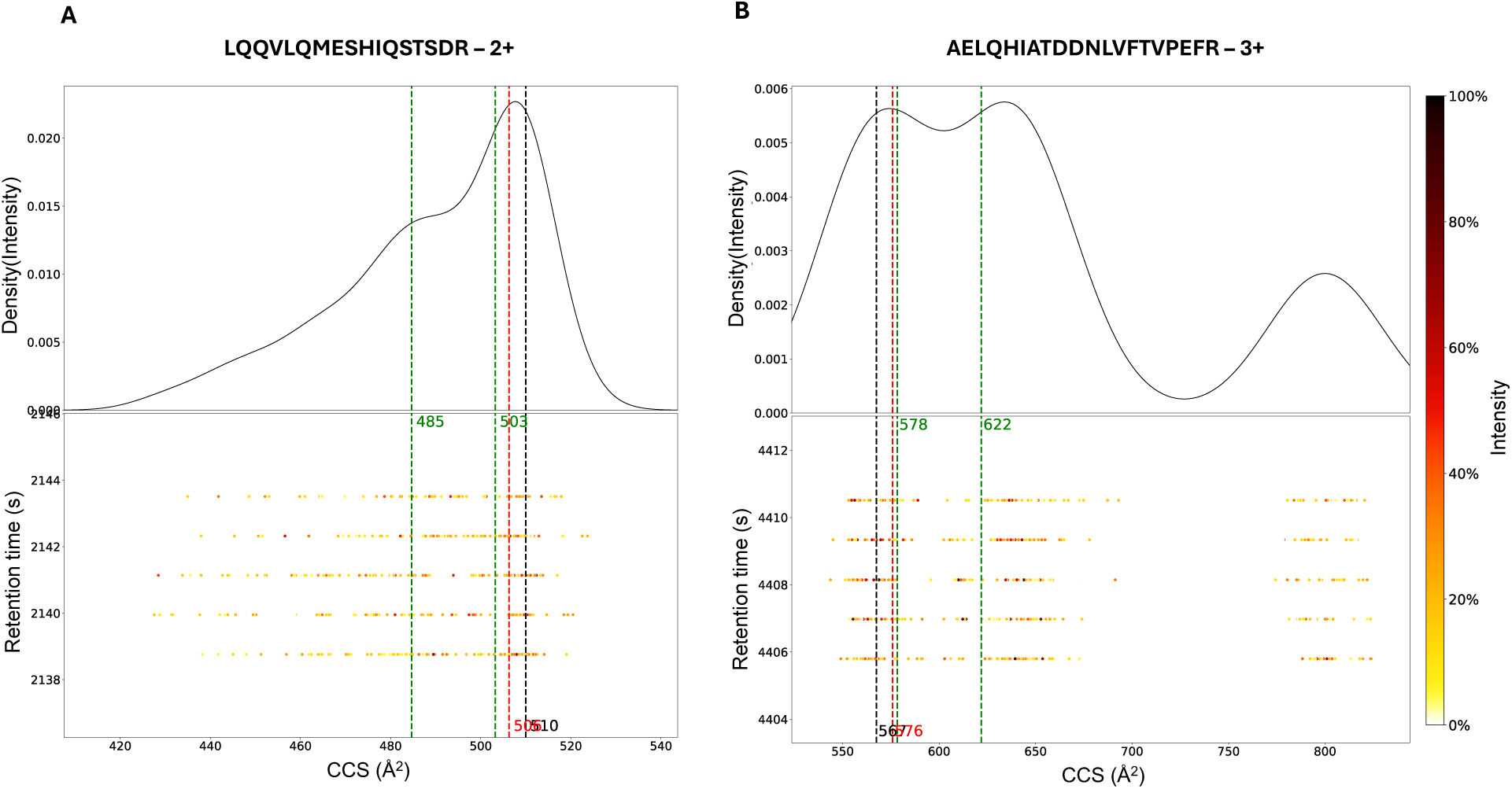
Mobilograms of selected precursor ions demonstrating remaining challenges in CCS prediction. A) This peptide ion (UniProt ID: Q14974) exhibits an ion mobilogram with unclear distinction between conformers, especially in the shoulder of the main conformation. B) Precursor ion (UniProt ID: P12111) clearly exhibiting three peaks in the ion mobility distribution. CCS value reported by Maxquant is indicated by a black dashed line. Classical IM2Deep prediction (single-output) indicated by red dashed line. CCS predictions made by multiconformational model indicated by green dashed lines.

Another intriguing aspect is the feasibility of predicting whether a precursor will exhibit multiconformational behavior. It might be the case that all precursors have the potential to exhibit multiple conformations depending on experimental conditions. Further investigations into the factors that influence multiconformationality are essential to facilitate the development of such predictors. Combining a predictor for multiconformational behavior with a multi-output CCS predictor can be a very effective tool for improved identification confidence.

However, larger challenges remain, particularly in distinguishing between closely related conformers due to overlapping ion mobility signals and minimal CCS differences (Figure 8A). Because search engines typically report apex CCS values, future research should focus on accurately linking ion mobility measurements to identified peptides. This would enable the prediction of entire ion mobility distributions rather than discrete CCS values. Accurately capturing these distributions would require advanced modeling techniques capable of handling the inherent complexity and variability in ion mobility data. Advanced deep learning techniques, such as variational autoencoders or generative adversarial networks that can integrate experimental information, could be leveraged to address these challenges. Continued data collection and sharing within the proteomics community are vital for the advancement of CCS (and other peptide behavior) prediction models and will facilitate the development of more sophisticated models by providing the necessary data diversity and volume.

The improvements provided by the multiconformational IM2Deep model have several implications for proteomics research. Improved CCS prediction accuracy can lead to more confident identification of peptides. Furthermore, the ability to predict CCS for multiple conformations can offer deeper insights into the structural dynamics and behavior of peptides in the gas phase.

## Conclusion

In this study, we enhanced IM2Deep by implementing a multi-output deep learning approach to predict collision cross sections for peptides exhibiting multiple conformations. Our findings demonstrate that the enhanced IM2Deep model accurately predicts CCS values for various conformers while maintaining high precision for peptides with a single observed conformation. These findings highlight the model’s potential to improve peptide identification and understanding of peptide behavior in the gas phase.

## Availability

IM2Deep is open source under the permissive Apache-2.0 license and is freely available as a Python package through PyPI and Bioconda. The source code is available on Github at https://github.com/compomics/IM2Deep.

IM2DeepTrainer, a Python package used to train new (multi-output) IM2Deep models is open source under the permissive Apache-2.0 license and is freely available as a Python package on PyPI. The source code is available on Github at https://github.com/rodvrees/IM2DeepTrainer.

All data and scripts required to reproduce the presented results are available on Zenodo at 10.5281/zenodo.14886715

## Author contributions

Robbe Devreese: Conceptualization, methodology, software, validation, formal analysis, visualization, investigation, writing – original draft

Alireza Nameni: Methodology, writing – review & editing

Arthur Declercq: writing – review & editing

Emmy Terryn: Investigation, data curation

Ralf Gabriels: Writing - review & editing

Francis Impens: Validation, writing - review & editing

Kris Gevaert: Validation, writing - review & editing

Lennart Martens: Conceptualization, supervision, funding acquisition, writing - review & editing

Robbin Bouwmeester: Conceptualization, methodology, software, validation, supervision, writing – review & editing

## Acknowledgements

R.D., A.D., R.G., L.M. and R.B. acknowledge funding from the Research Foundation Flanders (FWO) [1SH9O24N, 12B7123N, 1SE3724N, G010023N, G028821N, 12A6L24N]. A.N. acknowledges funding from the European Union’s Horizon 2020 research and innovation programme under the Marie Skłodowska-Curie grant agreement N° 956148. L.M. and F.I. acknowledge funding from the Horizon Europe Projects BAXERNA 2.0 [101080544] and COMBINE [101191739], L.M. acknowledges funding from the Ghent University Concerted Research Action [BOF21/GOA/033] and funding from the CHIST-ERA project ODEEP-EU [G0GDV23N].

## Supplementary Information

**Supplementary Table 1:**
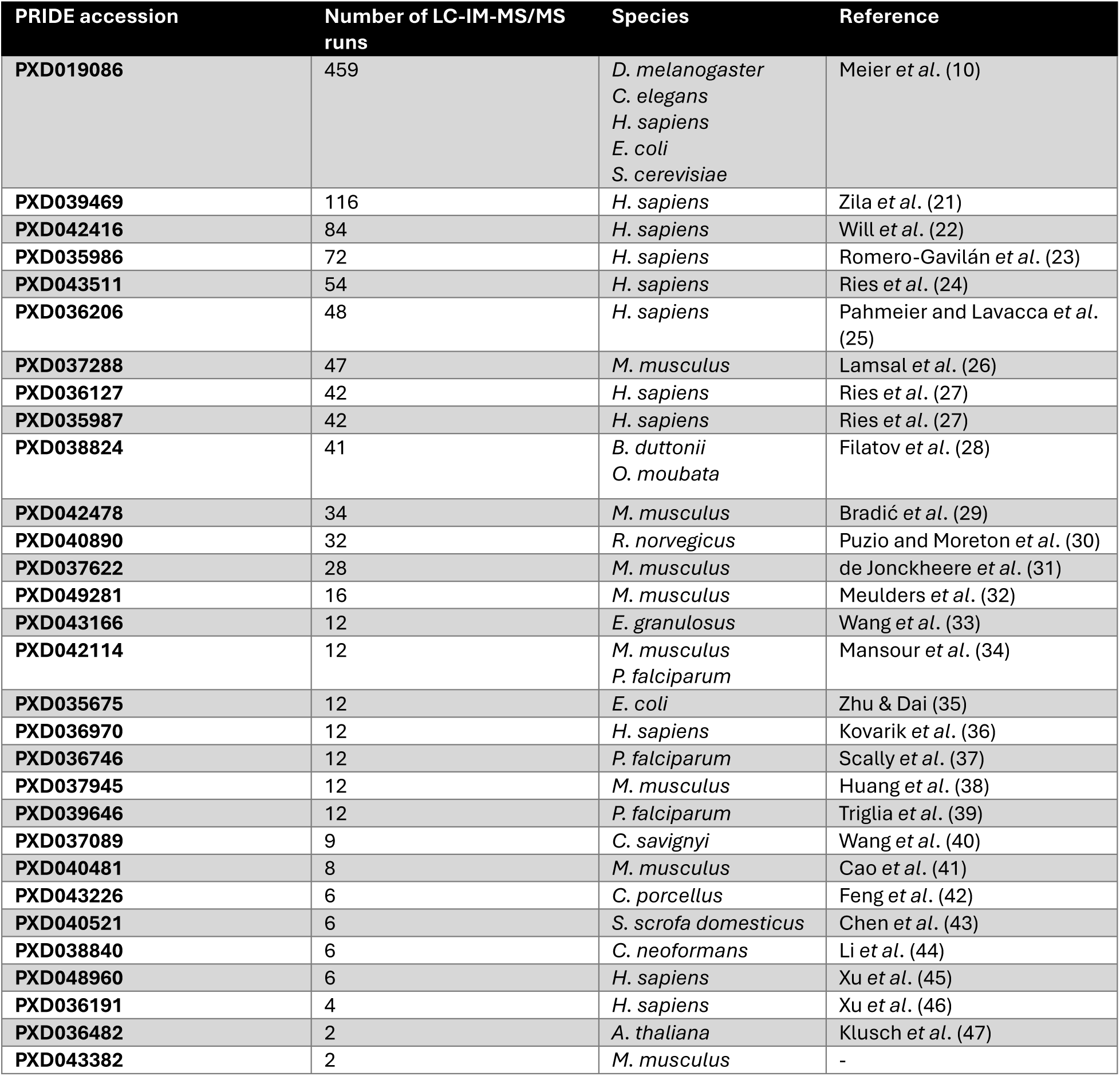
Overview of the public datasets used for further processing, along with the number of LC-IM-MS/MS runs they consist of and species from which data was acquired.

**Supplementary Figure 1:**
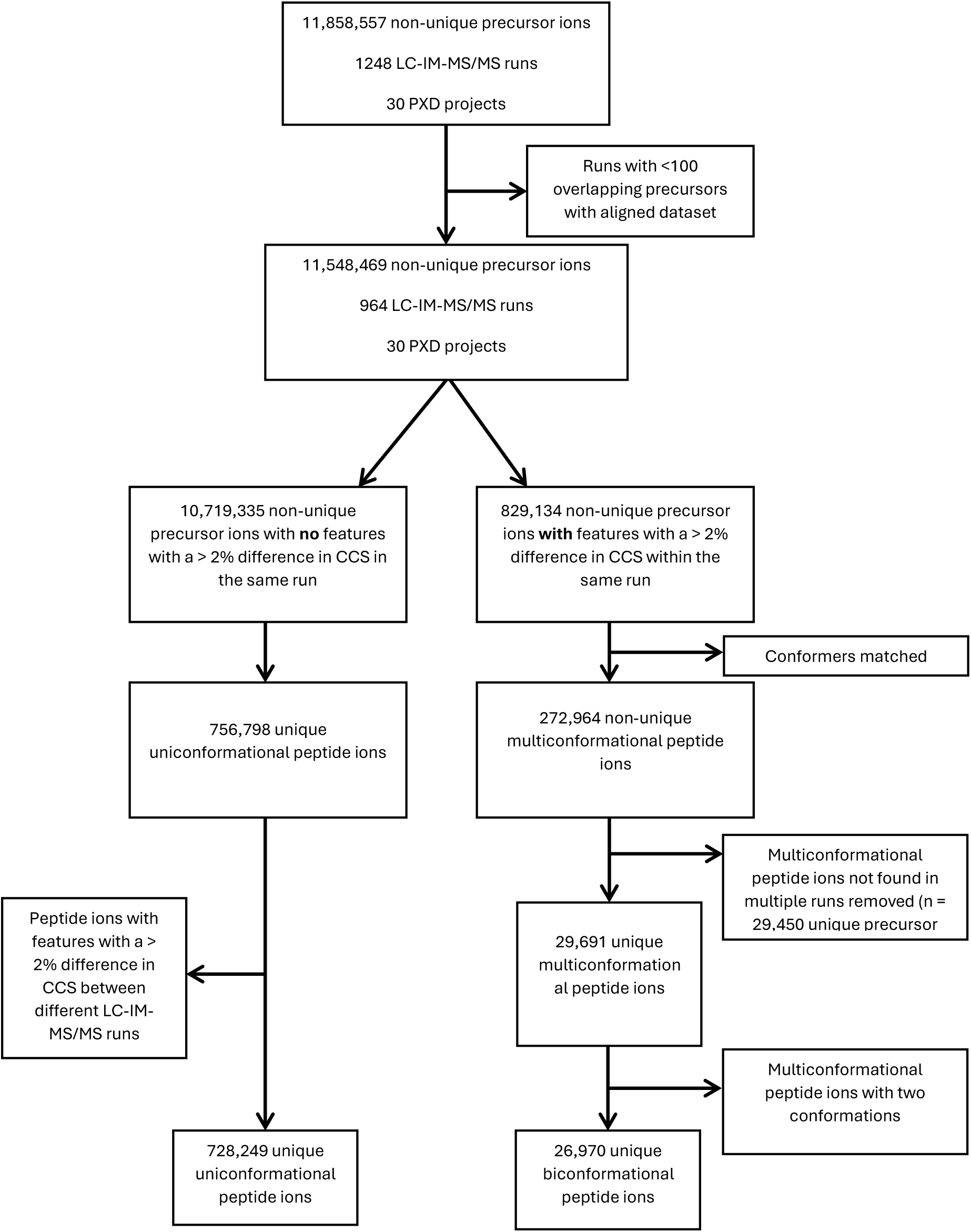
Processing workflow for the creation of the multiconformational precursor ion dataset.

**Supplementary Table 2:** Hyperparameters of the final multi-output IM2Deep model.

| Hyperparameter | Value |
| --- | --- |
| Batch size | 64 |
| Learning rate | 0.0001 |
| Activation function | Tanh for one-hot encoding path, Leaky ReLU for other paths |
| Leaky ReLU alpha | 0.1 |
| Maximum activation value | 20 |
| L1 lambda | 0.00001 |
| Max epochs | 500 |
| Kernel size | 4 for atom composition path, 2 for other paths |
| Number of filters | Atom composition path: 256<br>Diatom composition path: 128<br>One hot encoding path: 2 |
| Number of dense units | Global features path: 16<br>Concatenated path: 128 |
| Difference weight | 5 |

**Supplementary Figure 2:**
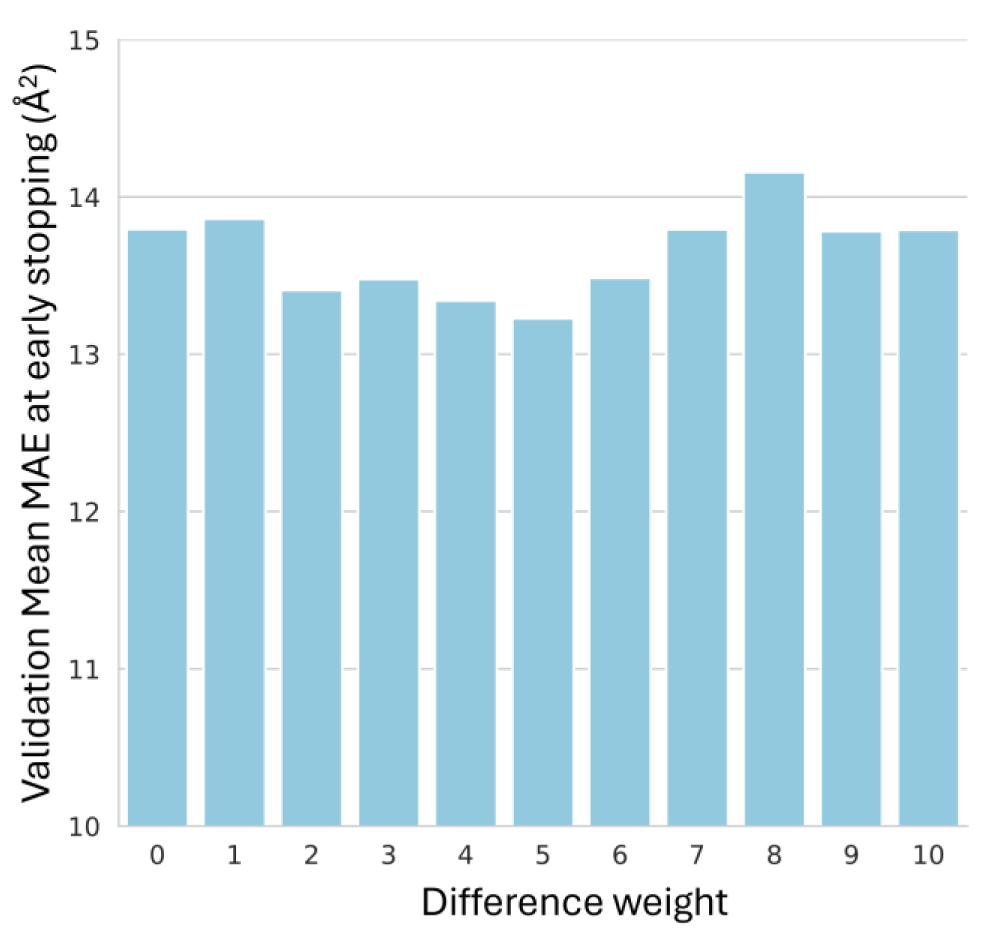
Validation accuracy for models trained with difference weights from 0 to 10.

**Supplementary Figure 3:**
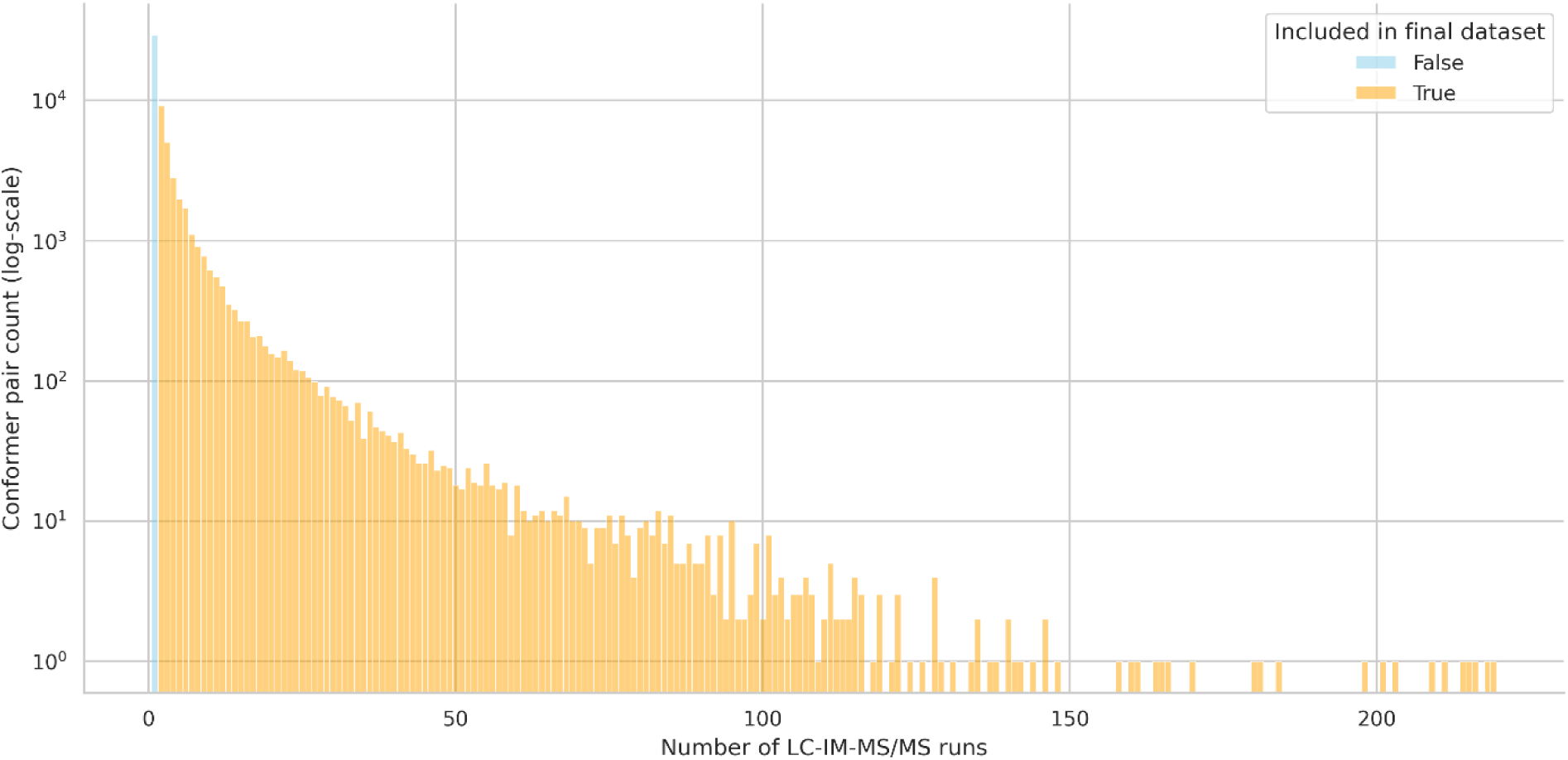
Distribution of conformer counts identified across multiple LC-IM-MS/MS runs, illustrating the impact of filtering on the final dataset. X-axis represents the number of runs in which a conformer pairing was identified, while the y-axis (log-scale) shows the total number of conformer pairs. Yellow bars represent conformers that were included in the final dataset after filtering.

**Supplementary Figure 4:**
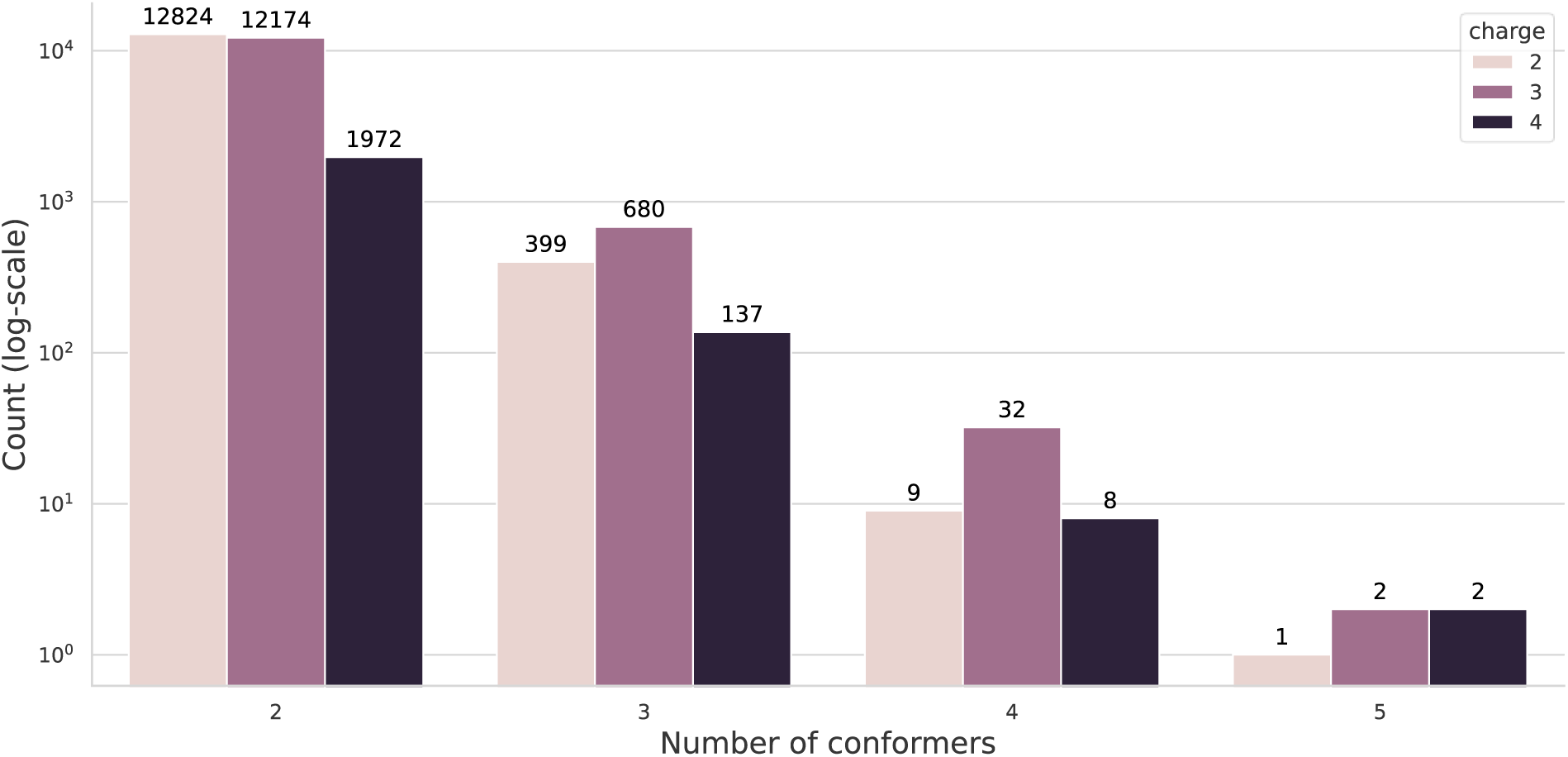
Distribution of the number of conformers observed per peptidoform-charge state pair in the multiconformer dataset after filtering, for each charge state. X-axis represents the number of conformers identified for each precursor, and the y-axis (log-scale) shows the count of peptidoform-charge state pairs.

**Supplementary Figure 5:**
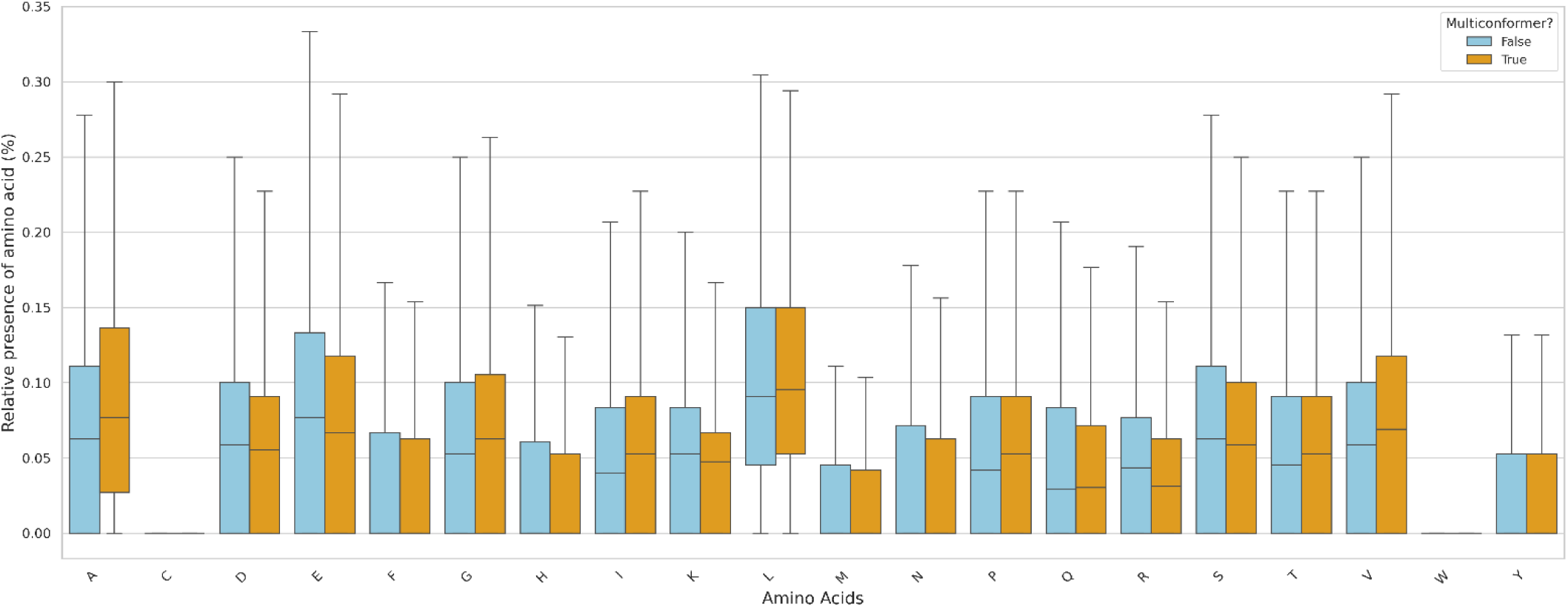
Boxplots comparing the relative presence of all 20 amino acids between multiconformational and uniconformational peptides.

**Supplementary Figure 6:**
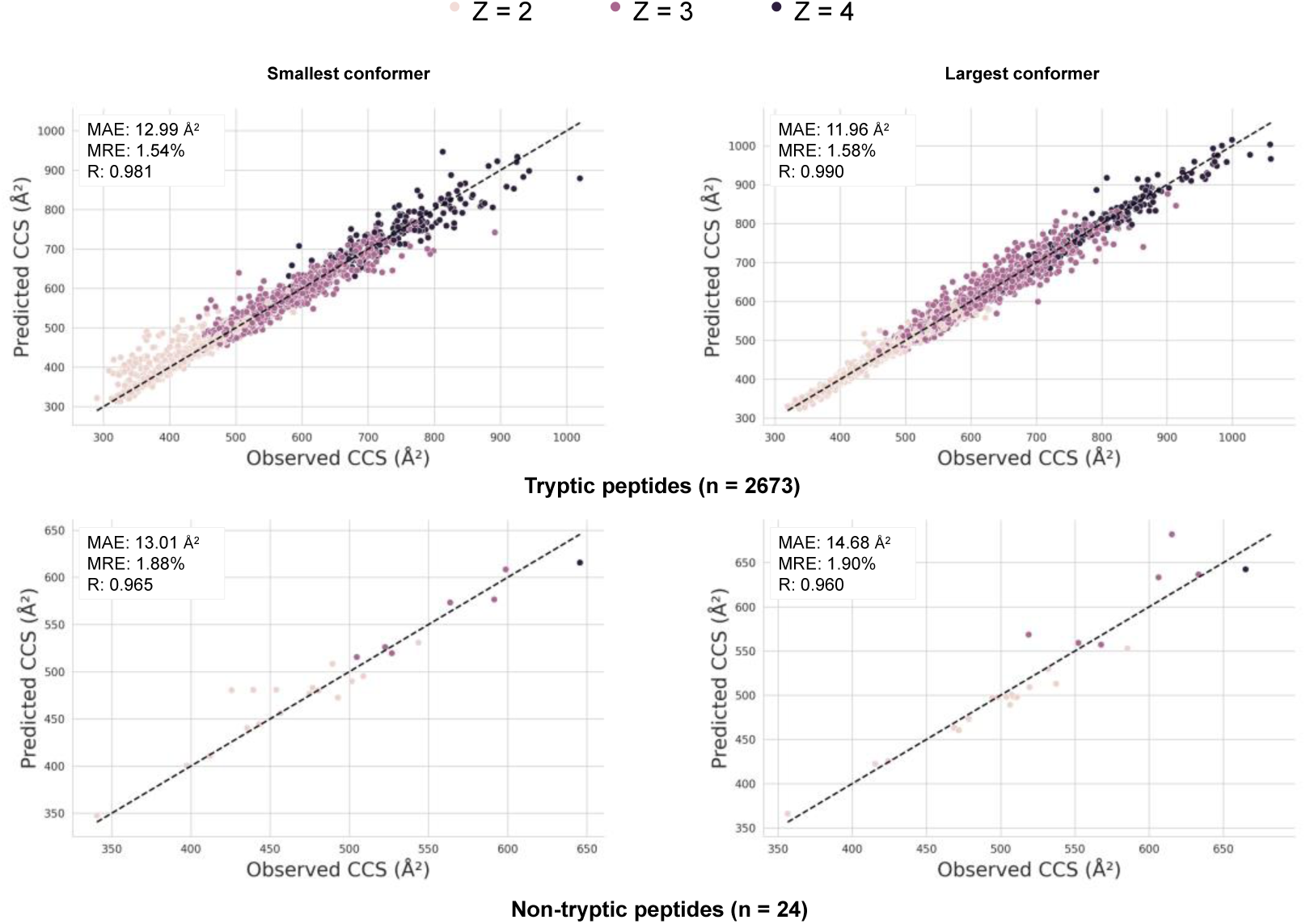
Scatter plots comparing predicted versus observed CCS values for the transfer-learned multi-output IM2Deep model, where performance on tryptic and non-tryptic precursor ions in the test set are plotted separately.

